# A type IV Autotaxin inhibitor ameliorates acute liver injury and non-alcoholic steatohepatitis in mice

**DOI:** 10.1101/2022.05.19.492626

**Authors:** Richell Booijink, Fernando Salgado-Polo, Craig Jamieson, Anastassis Perrakis, Ruchi Bansal

**Author notes:** Corresponding authors **Correspondence:** Anastassis Perrakis, NKI Dept B8, Plesmanlaan 121, 1066CX Amsterdam, The Netherlands, Ruchi Bansal, University of Twente, Drienerlolaan 5, 7522NB Enschede, The Netherlands. These authors contributed equally to this work (Anastassis Perrakis, Ruchi Bansal).

## Abstract

An important but rather underexplored pathway implicated in liver disease is the lysophosphatidic acid (LPA) signaling axis. LPA acts through G-protein coupled receptors inducing downstream signaling pathways related to cell proliferation, differentiation, and migration, and is predominantly produced by the extracellular phosphodiesterase, Autotaxin (ATX). ATX has gained significant attention lately with an impressive number of ATX inhibitors (type I-IV) reported. Here, we aim to evaluate the therapeutic potential of a (yet unexplored) type IV ATX inhibitor, Cpd17, in liver injury. In this study, we first confirmed the involvement of the ATX/LPA signaling axis in human and murine diseased livers. Thereafter, we evaluated the effects of Cpd17, in comparison with the classic type I ATX inhibitor PF8380, *in vitro*. While both inhibitors attenuated induced cell injury phenotypes as assessed using various assays and specific readout parameters in hepatocytes, macrophages, and hepatic stellate cells (HSCs), Cpd17 appeared more effective. This prompted us to characterize the mechanism of action of both inhibitors *in situ* and *in vitro* in macrophages and HSCs, demonstrating that Cpd17 was more potent in inhibiting relevant signaling pathways, namely RhoA-mediated cytoskeletal remodeling, and phosphorylation of MAPK/ERK and AKT/PKB. Finally, we investigated the therapeutic potential of Cpd17 in two liver disease mouse models, CCl_4_-induced acute liver injury and diet-induced non-alcoholic steatohepatitis. We demonstrate that Cpd17 has an excellent potential for reducing liver injury in both disease models *in vivo*. We conclude that ATX inhibition, by type IV inhibitor in particular, has an excellent potential for clinical application in liver diseases.

## Introduction

Liver diseases are an immense burden on today’s modern society. Globally, millions of people suffer from liver diseases, causing approximately 2 million deaths per year^1^. Among several etiological liver diseases drug-induced acute liver injury^2^ and non-alcoholic fatty liver disease (NAFLD) and its severe form non-alcoholic steatohepatitis (NASH) have been recognized as a major public health concern in Western countries^3, 4^.

Liver injury caused by different factors such as drug intoxication, excessive fat and/or alcohol intake, and hepatitis B/C viral infections, triggers hepatocyte injury and necrosis. This injury is accompanied by endoplasmic reticulum stress, inflammation through reactive oxygen species, and release of several pro-inflammatory cytokines and chemokines^5^. Ongoing inflammation induces the release of pro- fibrogenic mediators, mainly by resident and infiltrating macrophages^6^, causing quiescent hepatic stellate cells (HSCs) to transdifferentiate into highly proliferative and activated HSCs (or myofibroblasts). These activated HSCs initiate a wound-healing response that is characterized by the accumulation of an excessive extracellular matrix (ECM) at the injured site. Upon persistent liver injury, chronic inflammation, and ECM accumulation lead to liver fibrosis (scarring), cirrhosis (end-stage liver dysfunction)^5, 7^, and/or hepatocellular carcinoma (primary liver cancer)^8^.

Among several pathways that have been associated with liver diseases is the lysophosphatidic acid (LPA) signaling axis^9^. Besides liver diseases, the LPA signaling pathway is also involved in pathological conditions that include chronic inflammatory disorders, fibrotic diseases, and tumor progression^10, 11^. Recently, the ATX-LPA signaling axis has also been shown to impede anti-tumor immunity by suppressing chemotaxis and tumor infiltration of CD8+ T cells^12^. The ATX/LPA pathway also plays a role in a plethora of biological processes including neurogenesis, vascular homeostasis, skeletal development and remodeling, and lymphocyte homing^13-16^.

LPAs are physiologically occurring, structurally simple water-soluble glycerol lysophospholipids that differ in length, and the number of saturated and unsaturated bonds of their alkyl chain. Their activity is exerted via the activation of downstream signaling pathways through G-protein coupled receptors (GPCRs) specific for LPA. To date, six LPA receptors have been identified, that belong to the endothelial differentiation gene (EDG) family (LPA_1_-LPA_3_) and the P2Y family (LPA_4_-LPA_6_)^17^, and couple with different Gα subunits upon activation. LPA receptor activation can occur through Gα(i/o) and Gα(12/13) driven cascades via phosphatidylinositol 3-kinase (PI3K) and RhoA respectively, resulting in cell survival and cytoskeletal remodeling respectively^18, 19^.

The majority of LPA in the blood is synthesized from lysophosphatidylcholine (LPC) through hydrolysis of its choline moiety by Autotaxin (ATX), a member of the ectonucleotide pyrophosphatase phosphodiesterase family of enzymes (thus also known as ENPP2)^20^. ATX is a multi-functional and multi-domain protein that possesses enzymatic lysophospholipase D (lysoPLD) activity^21, 22^. Structurally, ATX has a catalytic phosphodiesterase (PDE) domain, which accommodates a tripartite binding site: a hydrophilic shallow groove adjacent to the catalytic site that harbors the glycerol moiety of the lysolipid substrate; a hydrophobic pocket that binds the acyl chain; and a tunnel leading to the other side of the PDE domain^22^. Given the emerging association with diseases, ATX has gained significant pharmacological and pharmacochemical attention (reviewed by Geraldo et al.,^23^), and an impressive number of ATX inhibitors has been reported^24^. The only inhibitor to reach phase 3 clinical trial (for idiopathic pulmonary fibrosis) has been the GLPG1690 molecule^25^, a so-called type IV inhibitor. Type IV inhibitors occupy the tunnel and the pocket of ATX, in contrast with type I inhibitors e.g., PF8380, which targets the catalytic site and the pocket^24^.

Elevated ATX levels were observed in patients with different etiological liver diseases that correlated with disease severity including non-alcoholic fatty liver disease (NAFLD)^26-31^. Moreover, hepatocyte- specific genetic deletion of ATX resulted in abrogated liver damage, inflammation and diminished fibrosis, and deregulated fatty acid metabolism thereby attenuating HCC development suggesting ATX as an attractive drug candidate in liver fibrosis and cancer^32^. ATX pharmacological inhibition in liver disease has been evaluated in only a few pre-clinical studies, with varying outcomes. The type I inhibitor PF8380, was found to reduce fibrosis, but not inflammation and necrosis, in a CCl_4_-induced liver fibrosis mouse model^32^. An ATX type III inhibitor, PAT-505, showed significant reduction of fibrosis score in two NASH mouse models, but no significant effect on inflammation, ballooning, or steatosis^33^. The ATX inhibitor EX_31, a tetrahydrocarboline derivative with unclear binding mode, did not show any effect on liver-related markers in a 10-week CCl_4_-induced liver fibrosis and 14-week choline-deficient amino acid-defined diet (CDAA)-induced NASH rat models^34^. Interestingly, type IV ATX inhibitors have not yet been explored in liver diseases.

We have recently shown that mechanistic differences between type I and type IV inhibitors, rather than inhibitor potency in preventing the enzymatic hydrolysis of LPC to LPA alone, can lead to different physiological activity^35^. Here, we set out to examine the effect of type IV inhibitor in liver disease models, as, despite the clear involvement of the ATX/LPA signaling pathway in liver diseases, inhibition of ATX has been underexplored in relevant liver disease models. We started by comparing the effects of our previously reported type IV Cpd17 inhibitor^36^ (not yet explored in liver diseases), with the classic type I inhibitor PF8380 *in vitro*. Cpd17, designed by fusing parts of type I and type III inhibitors, occupies the binding pocket and the tunnel but does not interact with the catalytic site. While both Cpd17 and PF8380 inhibitors possess similar potency in inhibiting LPC 18:0 to LPA conversion, our results demonstrate that Cpd17 exhibits higher potency in inhibiting downstream LPA signaling pathways *in situ* and *in vitro*. We were thus encouraged to examine Cpd17 in CCl_4_-induced acute liver injury, and methionine- and choline-deficient (MCD) diet-induced NASH mouse models.

## Materials and methods

### Transcriptomic analysis

For ATX (ENPP2) analysis in human tissues, the transcriptomic datasets of liver tissue from patients with liver cirrhosis (GSE14323), and non-alcoholic steatohepatitis (GSE63067), from the National Center of Biotechnology Information Gene Expression Omnibus database (NCBI-GEO) were analyzed using GEO2R.

### Cell Lines

Human hepatic stellate cells (LX2 cells), provided by Prof. Scott Friedman (Mount Sinai Hospital, New York, NY, USA), were cultured in DMEM-Glutamax medium (Invitrogen, Carlsbad, CA, USA) supplemented with 10% fetal bovine serum (FBS, Lonza, Verviers, Belgium) and antibiotics (50U/mL Penicillin and 50μg/mL streptomycin, Sigma, St. Louis, MO, USA). Murine RAW 264.7 macrophages and human THP1 monocytes, obtained from the American Type Culture Collection (ATCC, Manassas, VA, USA), were cultured in Roswell Park Memorial Institute (RPMI) 1640 medium (Lonza) supplemented with 10% FBS (Lonza), 2mM L-glutamine (Sigma) and antibiotics. Human HepG2 cells (ATCC) were cultured in DMEM-high glucose medium (Lonza) supplemented with 10% FBS and antibiotics.

### AKT and ERK phosphorylation by Western blotting

100,000 LX2 or 20,000 RAW cells were seeded in 6- and 12-well tissue culture plates and allowed to grow for 24h in DMEM (GIBCO, Life Technologies) containing 10% FBS and antibiotics, containing 5ng/mL TGFβ (for LX2 cells), or 100ng/mL LPS and 10ng/mL IFNγ (for RAW cells). Next, they were washed twice with PBS and serum starved overnight. Mixes with 1μM 18:1 LPC, 1μM 18:1 LPA, 20nM ATX, 1μM PF8380, 1μM Cpd17, or 1μM Ki16425 were incubated for 30min in serum-free medium containing 0.05% (w/v) fatty acid-free BSA (total volume 1mL). Medium from the plates was removed and replaced with 1mL of ATX-inhibitor mixture. Cells were stimulated for 10min, medium was removed, and plates were immediately frozen on dry ice and stored at −80°C. For Western blotting, cells were lysed in RIPA buffer, supplemented with protease inhibitors (Pierce), 20mM NaF and 1mM Orthovanadate, and spun down. Protein concentration was measured using a BCA protein assay kit (Pierce). LDS sample buffer (NuPAGE, Invitrogen) and 1Mm dithiothreitol (DTT) were added to the lysate. 20μg total protein were loaded on SDS-PAGE precast gradient gels (4–12% Nu-Page Bis-Tris, Invitrogen), followed by transfer to nitrocellulose membrane. Non-specific protein binding was blocked by 5% BSA in PBS-Tween (0.1%); primary antibodies (Supplementary Table 1). They were incubated overnight at 4°C in PBS-tween with 5% BSA, containing 0.1% NaN_3_. Blots were then incubated for 1h at RT with monoclonal anti-β-actin antibody prepared in PBS-tween with 5% skimmed milk containing 0.1% NaN_3_. Horseradish peroxidase-conjugated secondary antibodies (Supplementary Table 1) were incubated for 1h at room temperature in PBS-tween with 2.5% BSA and developed using ECL Western blot reagent (Thermo Scientific).

### Rho GTPase Biosensor

A FRET pair consisting of RhoA-Cerulean3 and PKN fused to circularly permuted Venus was used^37^. The CRIB domain of PAK and HR1 region of PKN were used as the effector domain for activated Rac1/Cdc42 and RhoA, respectively. Experiments were performed in HEPES-buffered saline (containing 140mM NaCl, 5mM KCl, 1mM MgCl_2_, 1mM CaCl_2_, 10mM glucose, 10mM HEPES), pH 7.2, at 37°C. Cells were allowed to adhere overnight on uncoated coverslips, after which they were serum-starved and transfected with the indicated biosensor for 24h. Next, the coverslips were placed on a thermostatted (37°C) inverted Nikon Diaphot microscope and excited at 425nm. Donor and acceptor emission were detected simultaneously with two photomultipliers, using a 505nm beam splitter and optical filters: 470 ± 20nm (CFP channel) and 530 ± 25nm (YFP channel). The emission data were analyzed using the Fiji software and normalized to control cells. At least three independent experiments were analyzed for every condition (10 fields of view/condition, 3–5 cells/field of view, >30 cells/condition). FRET was expressed as the ratio between acceptor and donor signals, set at 1 at the onset of the experiment.

### Surface biotin labeling

For LPA1-HA internalization, LX2 cells were grown on 6-well plates 24h, medium was overnight in serum-free DMEM medium, LPA1-HA was overexpressed in LX2 cells upon transfection with Fugene6 at a 1:3 (μg/mL) ratio overnight. Between each of the previous steps, cells were washed twice with PBS. For LPA1-HA internalization, cells were stimulated with the indicated reagents serum-free DMEM containing 0.1% fatty acid-free BSA for 15min. Next, the cells were transferred to ice, washed in ice-cold PBS, and surface labeled with 0.2mg/mL Sulfo-NHS-SS-Biotin (Thermo Scientific) for 30min. Biotin-labeled LPA1-HA was detected using streptavidin beads (Pierce) and conjugated anti-HA antibody (3F10 from Roche Diagnostics; 1:1000).

### Effect of ATX inhibitors on lipid biogenesis in human hepatocytes (HepG2 cells)

HepG2 cells were treated with palmitate (0.2mM) in complete culture medium, supplemented with 1% BSA, with and without PF8380 or Cpd17 (1μM) for 48h. Thereafter, cells were washed twice with PBS, and either lysed with RNA lysis buffer or fixed and stained using an Oil-red-O staining kit (Sigma) as per manufacturer’s instructions. Experiments were performed as three individual experiments.

### Effects of ATX inhibitors on macrophages

Mouse RAW macrophages or human THP1 monocytes (with 50ng/mL phorbol 12-myristate 13-acetate, PMA, Sigma) were plated in 12-well plates (5×10^5^ cells/mL) and incubated overnight. They were then incubated with cell medium alone, M1 stimulus (100ng/mL LPS and 10ng/mL IFNγ) and M2 stimulus (10ng/mL IL-13 and 10ng/mL IL-4), with or without 1μM of PF8380 or Cpd17 for 24h. For wound healing analyses, a 200μL pipet was used to make a scratch on the surface of the cells. Pictures were taken at 0h and 24h after incubation with transmitted light microscopy. Afterwards, cells were lysed with an RNA lysis buffer for quantitative real-time PCR analyses. Experiments were performed as three individual experiments.

### The effect of ATX inhibitor Cpd17 on human LX2 cells

Cells were seeded in 12-well plates (1×10^5^ cells/ml) or 24-well plates (6×10^4^ cells/ml) and cultured overnight. To study the effect of the inhibitor, cells were starved with serum free medium overnight, and then treated with medium alone or 5ng/ml human recombinant TGFβ with or without 1μM PF8380 or Cpd17. A 200μL pipet tip was used to create a scratch in the cell monolayer (12-well plates). Images were taken at 0h and 24h after the scratch using transmitted light microscopy (Evos microscope). The percentage of cells migrated into the wound was measured by the difference in scratch diameter between 0 and 24h, normalized to untreated (control) cells. Thereafter, cells were lysed with RNA lysis buffer to perform quantitative real-time PCR analysis. In addition, cells (24 well plates) were fixed with acetone: methanol (1:1) solution, dried and stained for collagen I and α-SMA. Experiments were performed as three individual experiments.

### 3D-collagen I contraction assay

3D-collagen I gel contraction assay has been done as described earlier^38^. Briefly, a 5.0mL collagen suspension with 3.0mL collagen G1 (5mg/mL, Matrix biosciences, Morlenbach, Germany), 0.5mL 10x M199 medium (Sigma), 85μL 1N NaOH (Sigma) and 1.415mL of sterile water was prepared. This suspension was mixed with 1mL (2×10^6^) LX2 cells. 0.6 mL of the collagen-gel suspension was plated in a 24 well plate and incubated at 37°C/5%CO2 for 1h. Polymerized gels were then incubated with serum-free medium with or without TGFβ (5ng/mL) together with 1μM of PF8380 or Cpd17. Thereupon gels were detached from the bottom of the culture wells. Pictures were taken with a digital camera at 0h and 24h. The area of the gel was digitally measured and normalized to the area of the inner size of the well in each image. Three individual experiments were performed.

### Animal Experiments

All the animal experiments performed in this study were in accordance with the guidelines and regulations for the Care and Use of Laboratory Animals, Utrecht University, The Netherlands. The protocols were approved by the Institutional Animal Ethics Committee of the University of Twente, The Netherlands. 6- to 8-week-old male C57BL/6NRj mice purchased from Janvier Inc. Labs (Le Genest, St. Isle, France), were housed in standard SPF conditions in a Tecniplast IVC system (cage type 2L, blue line). For the carbon tetrachloride (CCl_4_)-induced acute liver injury model, male C57BL/6NRj mice were treated with a single intraperitoneal injection of olive oil (Sigma) or CCl_4_ (Sigma, 0.5ml/kg in olive oil) at day 1. At day 2 and day 3, CCl_4_-treated mice received intraperitoneal administration of 5mg/kg Cpd17 or vehicle treatment (n=5 per group). On day 4, all mice were sacrificed, blood and livers were collected for subsequent analysis. For the methionine and choline deficient (MCD) diet-induced NASH model, C57BL/6NRj mice were fed with a standard chow or MCD diet for 7 weeks. Starting at week 5, the mice received intraperitoneal administrations of 5mg/kg Cpd17 or vehicle treatment 2 times per week, for a duration of 3 weeks, n=6 per group. All mice were sacrificed 24h after the last treatment, blood and livers were collected for further analysis. Alanine aminotransferase (ALT), aspartate aminotransferase (AST), total plasma cholesterol levels and plasma triglycerides levels were measured by standard automated laboratory methods.

### Hydroxyproline assay

Total liver hydroxyproline concentration was determined using a hydroxyproline assay (Sigma), a commonly used method for quantifying total collagen content, according to manufacturer’s instructions. In short, 10mg liver tissue was homogenized in 100μL MilliQ, with a mechanical homogenizer (Ultra-Turrax, IKA), and transferred to a pressure-tight polypropylene vial with PTFE-lined cap. 100μL concentrated hydrochloric acid (12M) was added, the vials capped tightly, and samples were hydrolyzed at 95°C for 24h. Thereafter, samples were centrifuged at 10,000 x g for 3min. 50μL of supernatant was transferred to a 96 well plate, which was allowed to dry at 60°C overnight. To each well, 100μL of Chloramine T/Oxidation buffer mixture was added and incubated for 5min. Thereafter, 100μL of p-Dimethylaminobenzaldehyde (DMAB; Ehrlich’s Reagent) diluted with perchloric acid/isopropanol solution was added to each well and incubated for 90min at 60°C. Absorbance was measured at 560nm and results quantified using known standard.

### Immunohistochemistry

Liver tissues were harvested and transferred to Tissue-Tek optimum-cutting temperature (O.C.T) embedding medium (Sakura Finetek, Torrance, CA, USA), and snap-frozen in 1- methyl butane chilled on dry ice. Cryosections (7μm) were cut using a Leica CM 3050 cryostat (Leica Microsystems, Nussloch, Germany). The sections were air-dried and fixed with acetone for 20 min. Cells or tissue cryosections were rehydrated with PBS and incubated with the primary antibody (Supplementary table 2) for 1h at room temperature or at 4°C overnight. Formalin-fixed paraffin embedded (FFPE) sections were deparaffinized in xylene, and rehydrated in graded ethanol, and distilled water. Antigen retrieval was achieved by overnight incubation at 60°C in 0.1M Tris/HCl buffer (pH 9.0) followed by the overnight incubation with the primary antibody. For tissue sections, endogenous peroxidase activity was blocked by 3% H_2_O_2_ prepared in methanol for 30min. Cells or sections were then incubated with horseradish peroxidase (HRP)-conjugated secondary antibody for 1h at room temperature and thereafter incubated with HRP-conjugated tertiary antibody for 1h at room temperature. Thereafter, peroxidase activity was developed either using AEC (3-amino- 9ethylcarbazole) substrate kit (Life Technologies, Carlsbad, CA, USA) for the cryosections or DAB (3,3’- Diaminobenzidine) substrate kit (Pierce, Thermo Scientific) for the FFPE sections for 20min, and nuclei were counterstained with hematoxylin (Sigma). Cells or sections were mounted in Aquatex mounting medium (Merck, Sigma) for cryosections or VectaMount (VectorLabs) for FFPE sections following dehydration in graded ethanol. The staining was visualized, and images were captured using light microscopy (Nikon eclipse E600 microscope, Nikon, Tokyo, Japan). Furthermore, sections were scanned using Hamamatsu NanoZoomer Digital slide scanner 2.0HT (Hamamatsu Photonics, Bridgewater, NJ, USA).

### Immunofluorescence

Cells were fixed with acetone: methanol (1:1) and air-dried for 20min. The cells were rehydrated with cold PBS and incubated with collagen I primary antibody (1:100) for 1h at room temperature. Cells were then incubated with Alexa-488-labeled secondary antibody (1:100) for 1h at room temperature. Thereafter, the cells were fixed with DAPI-containing antifade mounting medium (Merck, Sigma). The staining was visualized, and images were captured using fluorescence microscopy (Evos).

### Oil-Red-O staining

Oil-Red-O stock solution was prepared by dissolving 0.3g Oil-Red-O (Sigma) in 100mL of isopropanol. Sections were fixed in 4% formalin for 20min and then stained with Oil-Red-O as per manufacturer’s instructions. Briefly, formalin-fixed sections were rinsed with 60% isopropanol followed by staining with freshly prepared Oil-Red-O working solution for 15min. Thereafter, sections were rinsed with 60% isopropanol and nuclei were counterstained with hematoxylin (Fluka Chemie, Sigma). Finally, sections were washed with tap water and mounted with Aquatex mounting medium (Merck, Sigma).

### Hematoxylin and Eosin staining

Sections were fixed with 4% formalin for 20min and then rinsed with distilled water. The sections were incubated with hematoxylin for 15min followed by washings with tap water. Thereafter, sections were incubated with eosin solution for 1.5min followed by washing in 96% ethanol, dehydration with ethanol and were mounted with VectaMount mounting medium (Vector Laboratories, Burlingame, CA).

### Quantitative histological analysis

For quantitative histological analysis, high resolution scans were viewed using NanoZoomer Digital Pathology (NDP2.0) viewer software (Hamamatsu Photonics. About 20 images (100x) of each entire section (from NDP) were imported into ImageJ and were analyzed quantitatively at a fixed threshold. All the primary antibodies used in this study have been pre-tested for specificity. The staining performed in the study included the negative control (without primary antibody) to confirm the specificity of the staining and showed no non-specific staining.

### RNA extraction, reverse transcription, and quantitative real time PCR

Total RNA from cells and liver tissues was isolated using GenElute Total RNA Miniprep Kit (Sigma) and SV total RNA isolation system (Promega Corporation, Madison, WI, USA), respectively, according to manufacturer’s instructions. The RNA concentration was quantified with a UV spectrophotometer (Nanodrop Technologies, Wilmington, DE, USA). Total RNA (1μg) was reverse transcribed using the iScript cDNA Synthesis Kit (Bio-Rad, Hercules, CA, USA). All primers were purchased from Sigma-Genosys (Haverhill, UK). Real-time PCR was performed using 2x SensiMix SYBR and Fluorescein Kit (Bioline, QT615-05, Luckenwalde, Germany), 20ng cDNA and pre-tested gene-specific primer sets (listed in Supplementary table 3). The cycling conditions for the BioRad CFX384 Real-Time PCR detection system was 95°C for 10min, 40 cycles of 95°C/15s, 72°C/15s, and 58°C/15s. Finally, cycle threshold (Ct) values were normalized to reference gene GAPDH or 18S and fold changes in expression were calculated using the 2-ΔΔCt method.

### Statistical analyses

All data is presented as the mean ± standard error of the mean (SEM). The graphs and statistical analyses were performed using GraphPad Prism version 9.1.2 (GraphPad Prism Software, Inc., La Jolla, CA, USA). Comparisons with control groups were analyzed using unpaired students’ t-test and/or multiple comparisons between different groups were performed by one-way analysis of variance (ANOVA) with Bonferroni post-hoc test. The differences were considered significant at p <0.05.

## Results

### Upregulation of Autotaxin in non-alcoholic steatohepatitis and liver cirrhosis

Upregulation of ATX in chronic liver diseases, including NASH and liver cirrhosis, has been previously observed in animal models, as well as in human patients^28, 32^. To confirm these findings, ATX (ENPP2) gene expression has been analyzed in human and mice livers. Analysis of mRNA expression levels revealed significantly elevated ENPP2/ATX expression in human NASH patients, compared to healthy control livers **(Fig. 1a)**. Similar upregulation of ATX gene expression was also observed in patients with liver cirrhosis **(Fig. 1a)**. Consistent with human data, ATX gene expression in mouse livers was upregulated in the MCD diet-induced NASH mouse model as well as in our CCl_4_-induced liver fibrosis mouse model **(Fig. 1b)**.

**Fig. 1:**
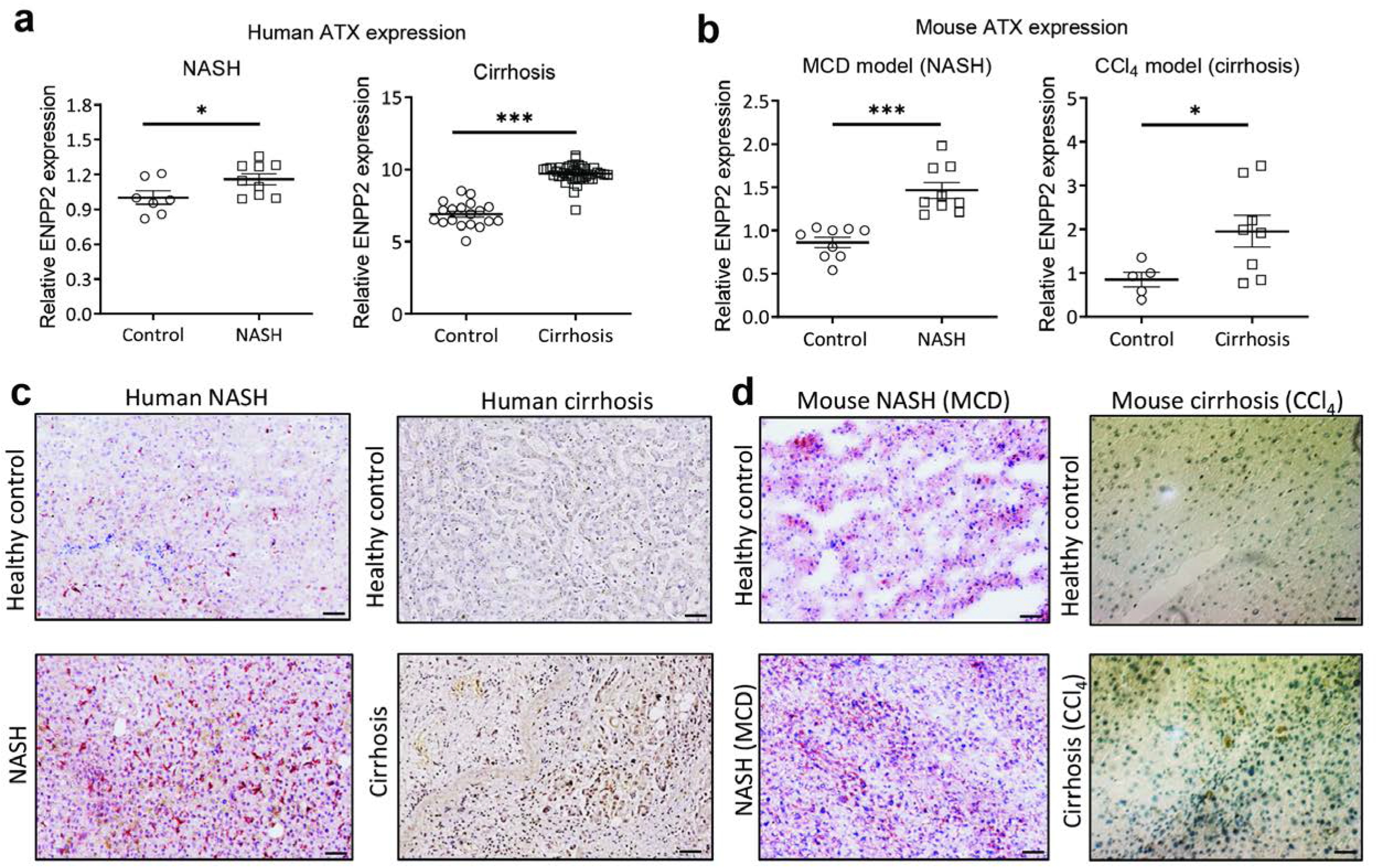
ATX (ENPP2) expression is upregulated in human and mouse livers. **(a)** ATX gene expression from publicly available datasets in NASH patients (n=9) compared to healthy controls (n=7); cirrhosis patients (n=40) compared to healthy controls (n=19). **(b)** Relative ATX gene expression in the MCD- diet-fed (n=9) compared to control mice (n=9); CCl_4_ (n=8) compared to control mice (n=5). **(c)** Representative images (scale=50μm) of ATX stained liver sections from human NASH compared to healthy controls; human cirrhosis compared to healthy controls. **(d)** Representative images (scale=50μm) of ATX stained liver sections (using AEC chromogen denoted by red color staining or DAB chromogen denoted by brown color staining; nuclei were stained blue with hematoxylin) from MCD- diet-fed NASH and CCl_4_-induced liver fibrosis mouse models compared to respective controls. Red color Mean ± SEM; two-tailed student’s t-test; *p<0.05, ***p<0.001 denotes significance versus respective healthy controls.

We further examined the protein expression of ATX in human and mouse livers by immunostaining. Hereby, we also found an increase in ATX protein expression in human NASH and cirrhotic livers compared to normal livers **(Fig. 1c)**. As expected, this was confirmed in both MCD-induced NASH and CCl_4_-induced cirrhosis mouse models compared to respective healthy controls **(Fig. 1d)**.

### Inhibition of ATX in hepatocytes and LPS/IFNγ-activated pro-inflammatory macrophages

To inhibit the function of ATX, and thus the production of LPA and the related downstream signaling, we focused on the well-characterized type I inhibitor PF8380 (**Fig. 2a**) and our type IV inhibitor Cpd17 (**Fig. 2b**). These two inhibitors possess similar potency in inhibiting the catalysis of LPC to LPA but distinct binding modes on the ATX tripartite binding site. Their effect on LPC species that we tested was similar (for 14:0, 16:0 and 18:1 LPC species) but showed interesting differences with Cpd17 being less potent in ameliorating hydrolysis of longer alkyl chain LPC species particularly 22:0 LPC (**Fig. 2 and supplementary Fig. 1**). We have thus decided to check both inhibitors in specific assays *in vitro* to evaluate their therapeutic potential.

**Fig. 2:**
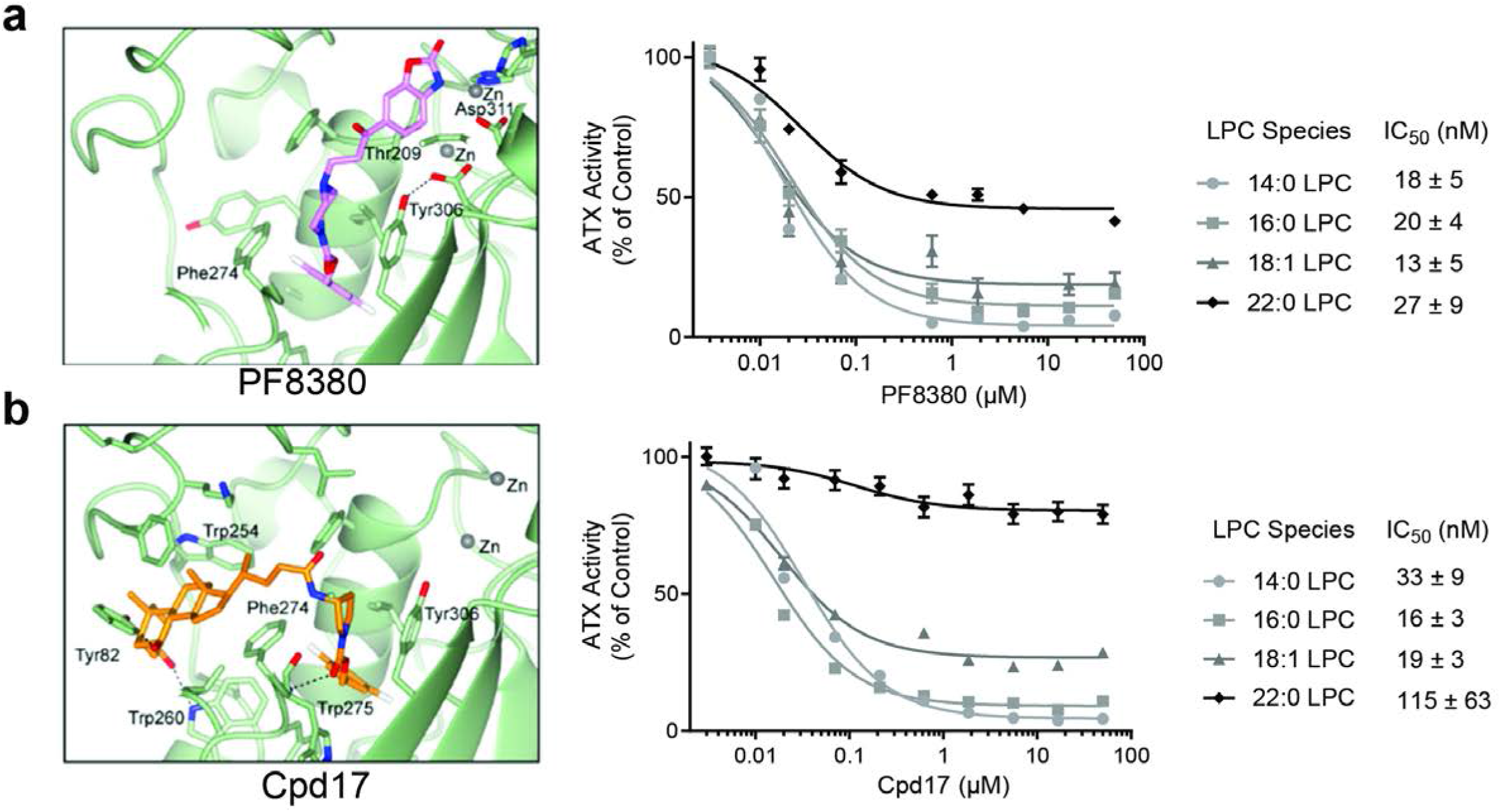
ATX inhibitors PF8380 and Cpd17 decrease LPA production. **(a)** Structural binding of PF8380 to ATX; Potency (IC_50_) of PF8380 for inhibiting the catalysis of LPC to LPA. **(b)** Structural binding of type IV inhibitor Cpd17 to ATX; Potency (IC_50_) of Cpd17 for inhibiting the catalysis of LPC to LPA.

Previously, it has been reported that different hepatotoxic stimuli stimulate hepatocyte ATX expression, leading to activation of fibrogenic pathways. Furthermore, hepatocyte-specific ablation and/or transgenic overexpression of ATX suggested a role of ATX/LPA in liver cirrhosis and HCC^32^. Moreover, since ATX has shown to be involved in fatty acid metabolism^32^, in this study, we first investigated the implication of pharmacological ATX inhibition using two different ATX inhibitors (Cpd17 and PF8380) in free fatty acid (Palmitate)-treated hepatocytes. To mimic NASH, we induced steatosis and hepatotoxicity to the human hepatoblastoma cell line, HepG2 cells, by exposing them to pathophysiological relevant concentrations of palmitic acid to mimic excessive influx of fatty acids into hepatocytes^39^ and found that ATX inhibition by Cpd17 significantly reduced lipid accumulation in hepatocytes as compared to PF8380 as assessed using Oil-red-O staining **(Fig. 3a)**.

**Fig. 3:**
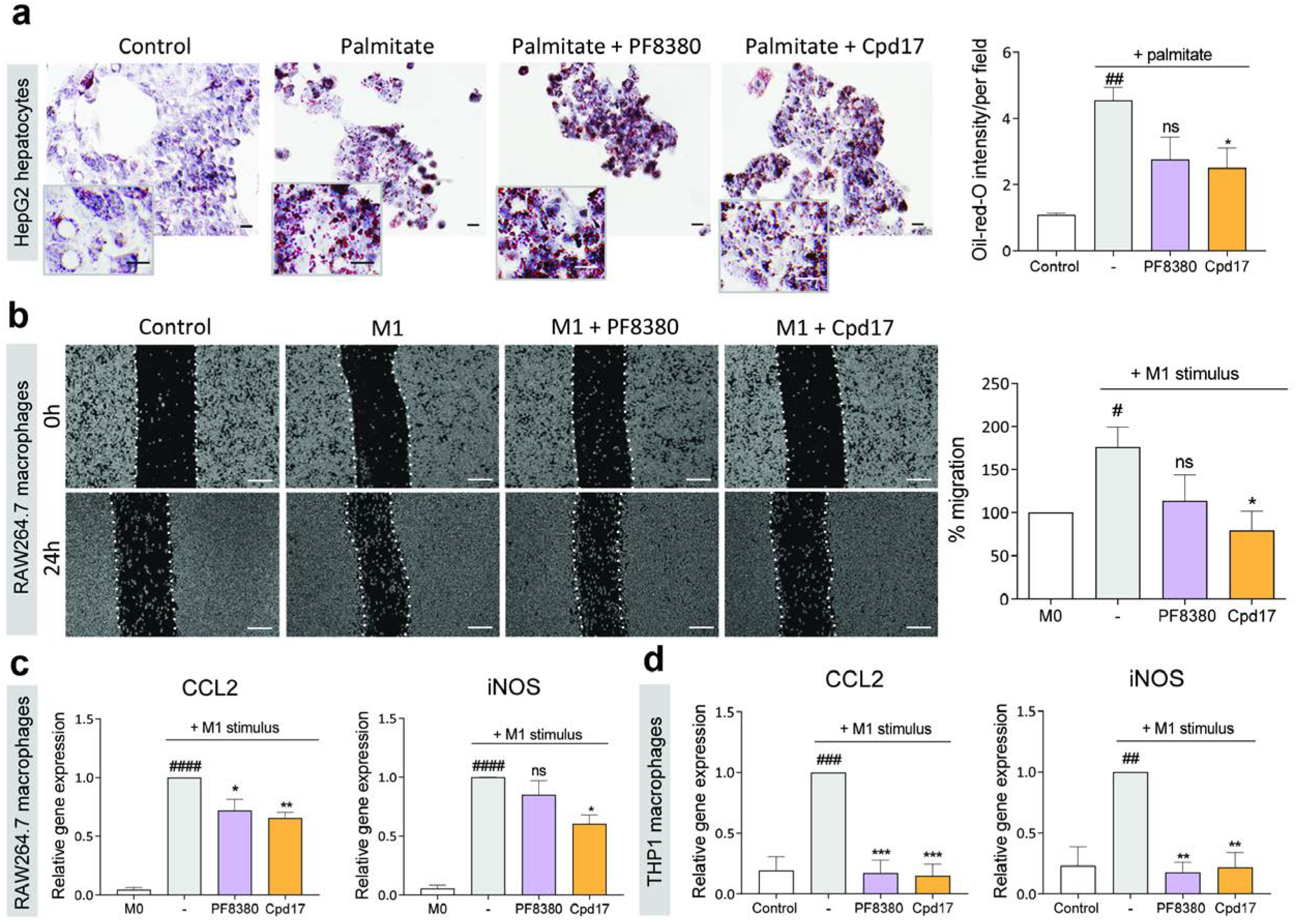
ATX inhibitors decrease steatosis in hepatocytes and pro-inflammatory expression and migration in LPS stimulated macrophages. **(a)** Representative images and quantification of HepG2 cells stained with oil-red-O after 48 hours treatment with or without 200μM palmitate with or without PF8380 or Cpd17 (scale=10μm). **(b)** Representative images (at 0h and 24h) (scale=50μm), and quantitative analysis (after 24h) of migration by control (M0) and LPS-induced M1 RAW 264.7 macrophages treated with medium alone, PF8380 (1μM) or Cpd17 (1μM). **(c)** Relative gene expression (normalized with GAPDH) for CCL2 and iNOS in control (M0) and LPS- and IFNγ- induced M1 RAW 264.7 macrophages treated with medium alone, PF8380 (1μM) or Cpd17 (1μM). **(d)** Relative gene expression (normalized with 18s RNA) for CCL2 and iNOS in control (M0) and LPS-induced PMA-treated human THP1 macrophages treated with medium alone, PF8380 (1μM) or Cpd17 (1μM). Mean + SEM; one-way ANOVA with Bonferroni post-hoc test; ^#^p<0.05, ^##^p<0.01, ^###^p<0.001, ^####^p<0.0001 denotes significance versus control (non-stimulated cells); *p<0.05, **p<0.01, ***p<0.001 denotes significance versus palmitate or M1 stimulated cells; ns=not significant.

Following hepatocellular damage, the inflammatory response plays a key role in the initiation and progression of liver fibrosis. Upon liver injury, infiltrating monocytes undergo differentiation into traditionally defined classically activated pro-inflammatory (M1) macrophages, which mediate the initial inflammatory response, while alternatively activated pro-resolving (M2) macrophages predominantly mediate the resolution phase^40^. We thus investigated the effect of the PF8380 and Cpd17 in classically activated mouse RAW macrophages. To investigate the effect of ATX inhibition on the migration of pro-inflammatory M1 (activated by IFN-γ and LPS) macrophages, we performed a scratch assay. 24h after making a scratch, LPS/IFNγ-activated pro-inflammatory macrophages showed a visibly significant increase in migration, compared to unpolarized (M0) macrophages **(Fig. 3b)**. Inhibition of ATX by PF8380 showed some decrease in macrophage migration; however, Cpd17 significantly reduced the percentage of cell migration compared to that of LPS/IFNγ-activated macrophages **(Fig. 3b)**. We further examined the efficacy of the inhibitors on the gene expression of inflammatory markers in both mouse and human macrophages. In mouse RAW264.7 macrophages, we observed that both PF8380 and Cpd17 significantly alleviated the LPS/IFNγ-induced expression of C-C chemokine 2 (CCL2), also known as monocyte chemotactic protein 1 (MCP1), one of the key chemokines that regulate migration and infiltration of monocytes/macrophages. Moreover, Cpd17, but not PF8380, also alleviated the expression of other inflammatory marker inducible nitric oxide synthase (iNOS) **(Fig. 3c)**. Similarly, in human THP1 macrophages, M1 activation through LPS and IFNγ led to an increase in gene expression of inflammatory markers CCL2 and iNOS. After 24h of treatment with both inhibitors, gene expression levels were significantly decreased **(Fig. 3d)**.

These results show that inhibition of ATX decreases lipid accumulation in hepatocytes and attenuates LPS/IFNγ-induced pro-inflammatory macrophage activation and migration, both in murine as well as human cells. These results confirm the importance of the LPA/ATX pathway in liver inflammation. Furthermore, the results suggest that Cpd17 is somewhat more efficient than PF8380 in ameliorating induced steatosis and inflammatory phenotype.

### Inhibition of ATX in TGFβ-activated hepatic stellate cells

The elevated ATX gene expression in patients could be recapitulated upon the activation of fibrogenesis *in vitro* in HSCs. This observation strengthened the previously reported association between ATX expression and the development of liver fibrosis and offered an attractive system for *in vitro* studies. We therefore investigated the effect of PF8380 and Cpd17 in TGFβ-activated LX2 cells (immortalized human HSCs). Upon TGFβ activation, immunostainings revealed an increase in fibrosis-related proteins, collagen I (the major ECM protein) and α-SMA (the HSCs activation marker) **(Fig. 4a)**; addition of PF8380 or Cpd17 visibly decreased the protein levels **(Fig. 4a)**. Correspondingly, gene expression levels of collagen I, α-SMA and platelet-derived growth factor beta receptor (PDGFβR, involved in HSC proliferation and survival), increased significantly upon TGFβ activation, showed some (non-significant) decrease after PF8380 treatment, but stronger (significant) reduction after Cpd17 treatment **(Fig. 4b)**. During fibrogenesis, increase in liver stiffness and excessive ECM accumulation is mediated by HSCs differentiation into myofibroblasts as well as HSCs migration to the injured site^41^.

**Fig. 4:**
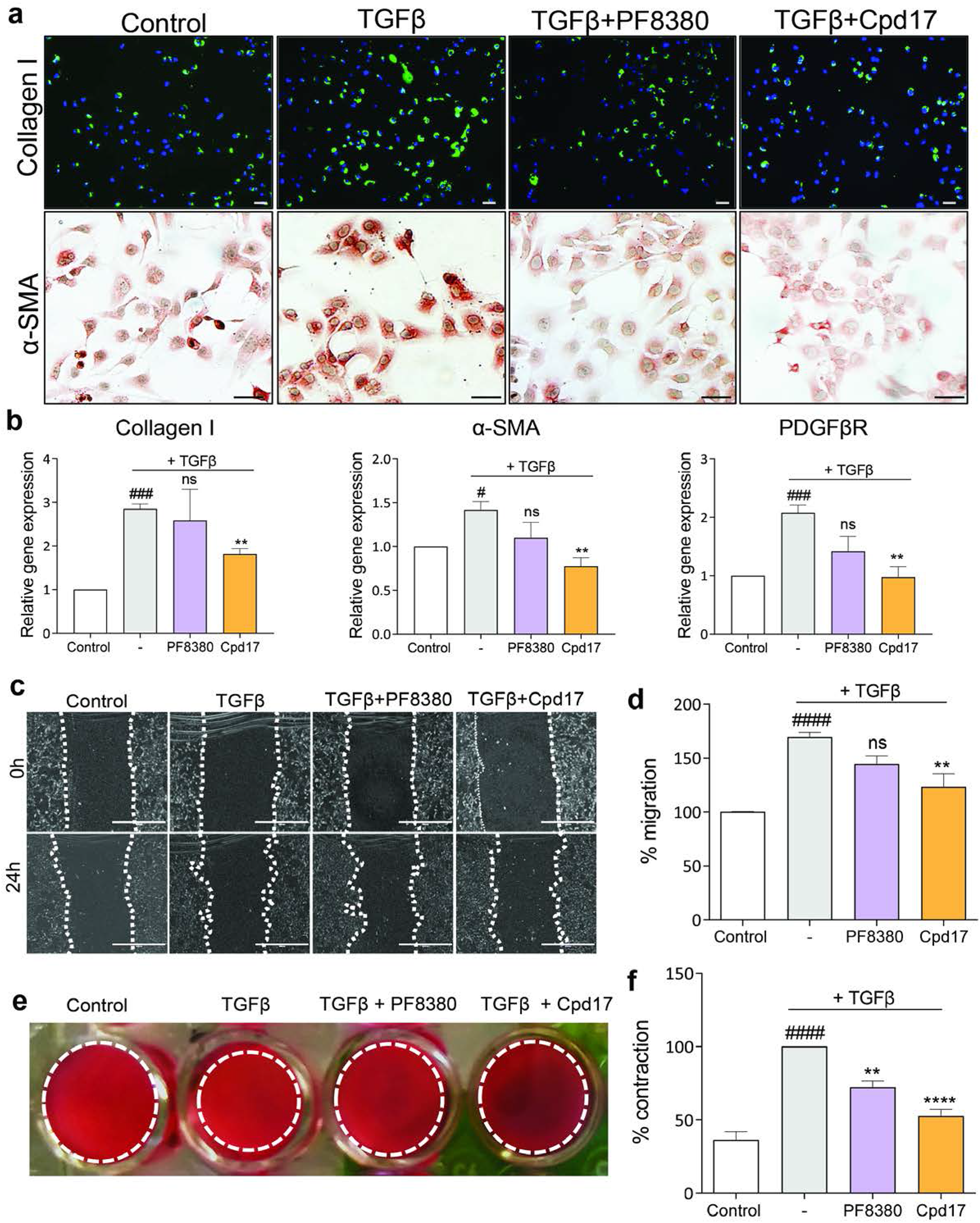
ATX inhibition decreases TGFβ-induced HSCs activation, migration, and contraction. **(a)** Collagen I and α-SMA stained images (scale=50μm) of control and TGFβ-activated LX2 cells with or without PF8380 (1μM) or Cpd17 (1μM) (n=3). **(b)** Relative Collagen I, α-SMA and PDGFβR gene expression (normalized with GAPDH) in control and TGFβ-activated LX2 cells with or without PF8380 (1μM) or Cpd17 (1μM) (n=3). **(c)** Representative images (at 0h and 24h) (scale=100μm) and **(d)** quantitative analysis (after 24h) of migration by control and TGFβ-activated LX2 cells with or without PF8380 (1μM) or Cpd17 (1μM) (n=3). **(e)** Representative images and **(f)** quantitative analysis showing 3D collagen gel contractility of control and TGFβ-activated LX2 cells with or without PF8380 (1μM) or Cpd17 (1μM) (n=3). Mean + SEM; one-way ANOVA with Bonferroni post-hoc test; ^#^p<0.05, ^###^p<0.001, ^####^p<0.0001 denotes significance versus control (non-stimulated cells); *p<0.05, **p<0.01, ***p<0.001 denotes significance versus (TGFβ) stimulated cells.

The migratory potential of activated HSCs was evaluated by a scratch assay. As observed previously, Cpd17 significantly reduced the wound healing response **(Fig. 4c-d)**, indicating a reduction in the migratory properties of the LX2 cells. Finally, we studied the effect of the two ATX inhibitors on the contractile properties of the LX2 cells using a 3D-collagen I contraction assay. Results showed that both PF8380 and Cpd17 significantly reduced TGFβ-induced collagen I contraction of LX2 cells after 24h of treatment **(Fig. 4e-f)**; consistent with other results the effect was more pronounced with Cpd17 treatment.

Overall, these results further solidify the significant role of the LPA/ATX pathway in TGFβ-induced activation, migration, and contractility of human HSCs, and confirm that the ATX type IV inhibitor Cpd17 is more efficient in reducing the fibrotic effects caused by HSC activation.

### PF8380 and Cpd17 differentially inhibit downstream signaling in LX2 cells and RAW macrophages

To investigate the mechanism by which PF8380 and Cpd17 affect inflammatory and fibrotic responses, we sought to dissect their downstream signaling effects in macrophages and HSCs, respectively. A short overview of LPA-induced signaling pathways is depicted in **Fig. 5a**. As all downstream signaling depends on the levels of ATX and the LPA_1-6_ receptors, we first evaluated their gene expression levels in TGFβ activated LX2 cells and in LPS/IFNγ-activated pro-inflammatory M1 macrophages (**Supplementary Fig. 2**). We observed that all the LPA receptors (except LPA_4_), as well as ENPP2, were upregulated in TGFβ-activated LX2 cells; and all LPA receptors (except LPA_2_ and LPA_5_), and ENPP2 were significantly increased in LPS/IFNγ-induced pro-inflammatory M1 macrophages, consistent with existing literature^32, 42-44^ (**Supplementary Fig. 2**).

**Fig. 5:**
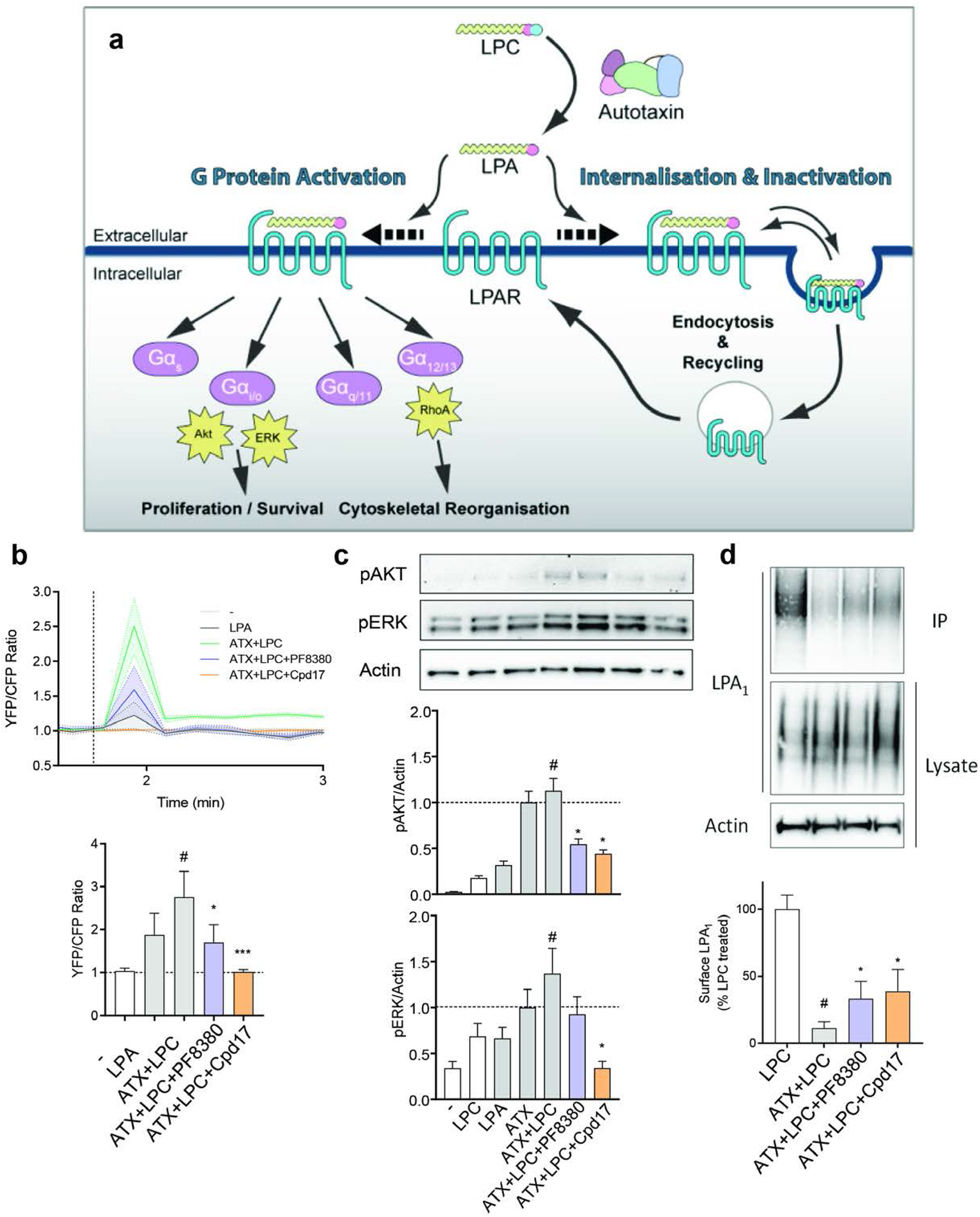
Inhibition of ATX results in blockade of LPAR-driven activation of stellate cells and macrophages. **(a)** Schematic overview of LPA-induced signaling pathways. **(b)** Top panel shows time-course stimulation to LPA and uninhibited or inhibited LPC-treated ATX followed by RhoA activation, measured as a YFP/CFP fluorescent ratio. Bottom panel depicts quantitation of the burst response to the stimulants. **(c)** Representative images and quantification showing Western-blot analysis for p-Akt and p-ERK normalized with β-actin in control, LPC, LPA, ATX, ATX+LPC, ATX+LPC+PF8380, and ATX+LPC+Cpd17 stimulated LX2 cells. **(d)** Representative images and quantification showing Western-blot analysis for LPAR1 on the cell surface and in lysate, normalized with β-actin in lysate, in LPC, ATX+LPC, ATX+LPC+PF8380, and ATX+LPC+Cpd17 stimulated LX2 cells. Mean ± SEM; one-way ANOVA with Bonferroni post-hoc test; ^#^p<0.05 denotes significance versus control (non-stimulated cells); *p<0.05, ***p<0.001 denotes significance versus ATX+LPC treated cells.

With respect to the signaling pathways, we first monitored the Gα(12/13)-mediated RhoA signaling pathway that leads to cytoskeletal reorganization, as it relates to HSCs morphology, migration and contractility^45^, which are hallmarks for mediating the fibrotic effect in HSCs. Consistently with the phenotypic observations, we observed that Cpd17, but not PF8380, strongly inhibited the ATX+LPC(LPA)-induced RhoA activation in LX2 cells (**Fig. 5b**). We then checked the Gα(i/o)-mediated phosphorylation of AKT and extracellular signal-regulated kinase (ERK). The maximal response in the amount of pAKT and pERK in LX2 cells is upon stimulation with ATX and the LPC substrate; this effect was ameliorated by both inhibitors in the case of pAKT accumulation, but only Cpd17 significantly suppressed pERK (**Fig. 5c**). We then checked the alternative signaling pathway involving receptor internalization, by biotin labeling of surface proteins, immunoprecipitation, and visualization of the amount of LPA_1_ in the cell surface (**Supplementary Fig. 3a**). While treatment with ATX and LPC significantly decreased the amount of LPA_1_ receptor in the cell surface, both inhibitors partially restored LPA_1_ surface levels (**Fig. 5d**). Finally, to confirm that the effects that we see are LPA-receptor dependent, we used the Ki16425 antagonist, mostly acting on LPA_1_, and demonstrated that its application had effects similar to those of the ATX inhibitors, and particularly Cpd17, both in LX2 cells as well as in LPS/IFNγ-differentiated M1 macrophages (**Supplementary Fig. 3b**).

These results demonstrate that ATX inhibition leads to signaling events on the downstream axis in cells that are relevant to the observed phenotypes. Crucially, the stronger effect of the Cpd17 inhibitor, specifically on RhoA activation, at least partially explains the consistent effect of the type IV inhibitor Cpd17 to ameliorate the effects in all our phenotypic assays. Our *in vitro* studies thus collectively suggested further analysis of the Cpd17 ATX inhibitor for therapeutic efficacy *in vivo*.

### Cpd17 decreases acute liver damage in CCl_4_-induced liver injury *in vivo*

The carbon tetrachloride (CCl_4_)-induced liver injury mouse model was our first choice for analyzing the *in vivo* efficacy of Cpd17 **(Fig. 6a)**. We observed that liver weights (normalized to body weight) were significantly increased in diseased mice compared to healthy control mice, and liver weights of the Cpd17-treated mice were significantly lower compared to the diseased mice **(Fig. 6b)**. Additionally, plasma transaminase (alanine transaminase, ALT) levels of mice treated with Cpd17 were also significantly reduced, compared to CCl_4_-treated mice **(Fig. 6c)**. Histological staining of the liver tissues showed an increase in inflammation, collagen I, and F4/80 protein expression in the CCl_4_-induced mouse model, as expected, Cpd17 notably decreased all these markers upon treatment **(Fig. 6d-e)**. Altogether, these results suggest that inhibition of ATX by Cpd17 treatment leads to reduced liver inflammation and fibrogenesis in a CCl_4_-induced liver injury mouse model.

**Fig. 6:**
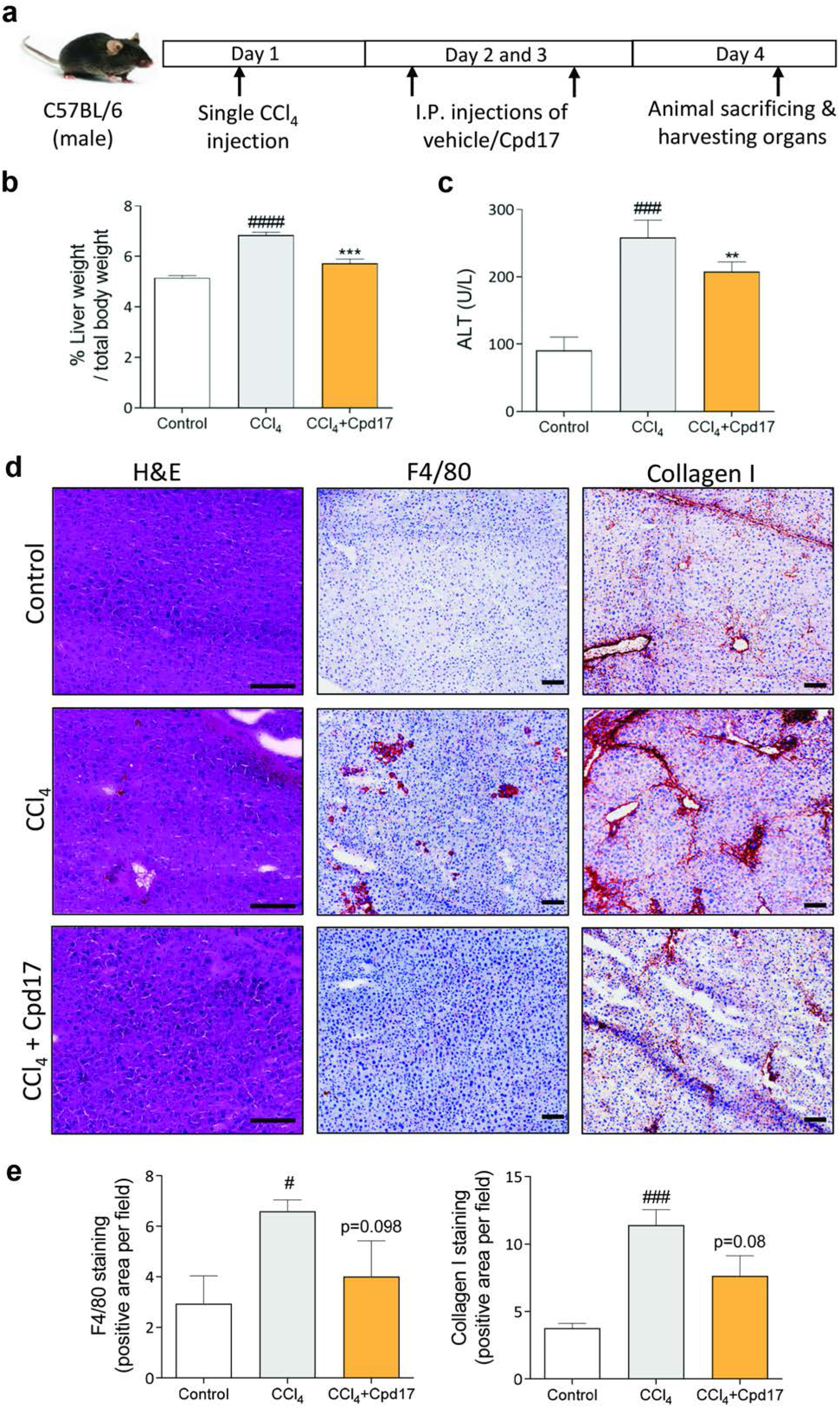
Cpd17 ameliorated CCl_4_ induced acute liver injury. **(a)** Schematic showing CCl_4_-mediated liver disease induction and Cpd17 treatment regimen. **(b)** Representative images (scale=100μm) of control (n=5), CCl_4_ (n=5) and CCl_4_+Cpd17 (n=4) liver sections stained with H&E, collagen-I and F4/80. **(c)** Liver weight to total body weight ratio. **(d)** Serum ALT levels. Mean ± SEM; one-way ANOVA with Bonferroni post-hoc test; ^###^p<0.001, ^####^p<0.0001 denotes significance versus controls; **p<0.01, ***p<0.001, ****p<0.0001 denotes significance versus CCl_4_ mice.

### Cpd17 ameliorates liver fibrogenesis and inflammation in methionine choline-deficient (MCD)-diet- induced NASH mouse model

Finally, we investigated the therapeutic efficacy of Cpd17 *in vivo* in a methionine- and choline-deficient (MCD)-diet induced NASH mouse model **(Fig. 7a)**. Morphological examination of the liver tissues directly after sacrificing, showed a distinctly visible pale and gross appearance suggesting hepatocellular damage in MCD-diet induced NASH mice as compared to the healthy control group; treatment with Cpd17 improved this appearance considerably **(Fig. 7b)**. Moreover, plasma ALT levels were significantly reduced in mice treated with Cpd17 compared to MCD-diet-fed NASH mice, further suggesting Cpd17 significantly improved liver function **(Fig. 7c)**. Total plasma cholesterol and triglyceride levels, indicators for liver steatosis (or NAFLD), increased significantly upon MCD diet- induced NASH were significantly decreased upon treatment with Cpd17 **(Fig. 7c)**.

**Fig. 7:**
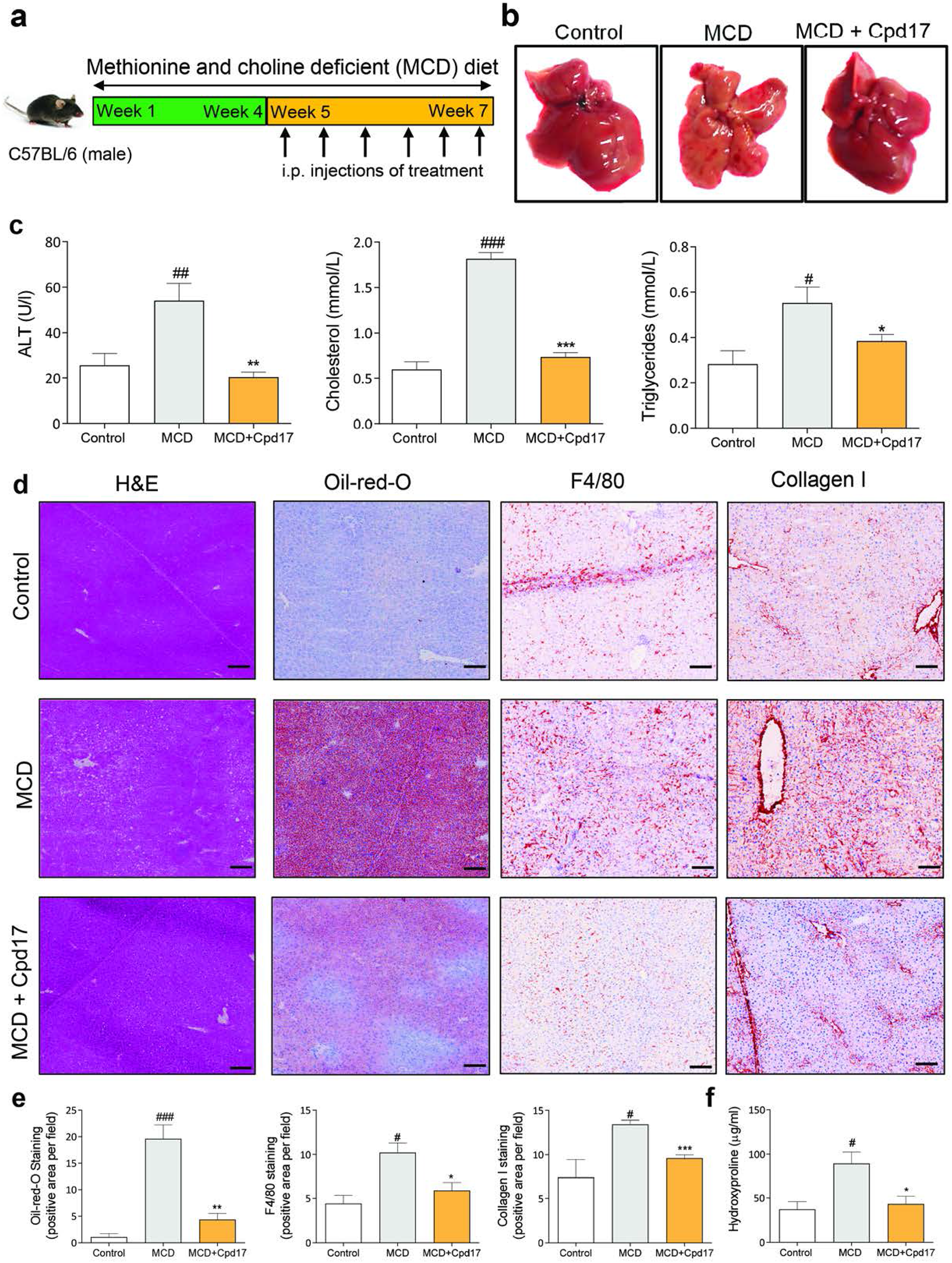
Cpd17 attenuated steatosis, inflammation and fibrosis in MCD-diet induced NASH mice. **(a)** Schematic showing NASH induction and Cpd17 treatment regimen. **(b)** Representative macroscopic images of livers from control (n=5), MCD (n=6) and MCD+Cpd17 (n=6). **(c-f)** Representative images (scale=100μm) and quantitative analysis of liver sections stained with **(c)** H&E, **(d)** oil-red-O, **(e)** F4/80 and **(f)** Collagen I. Mean + SEM; one-way ANOVA with Bonferroni post-hoc test; ^#^p<0.05, ^##^p<0.01, ^###^p<0.001 denotes significance versus controls; *p<0.05, **p<0.01, ***p<0.001 denotes significance versus MCD mice.

**Fig. 8:** ATX inhibition improved general liver disease levels in MCD-diet induced NASH mice. **(a-b)** Serum AST and ALT, **(c)** liver hydroxyproline, **(d)** serum cholesterol and **(e)** serum triglyceride levels in control (n=5), MCD (n=6) and MCD+Cpd17 (n=6) mice. Mean ± SEM; one-way ANOVA with Bonferroni post-hoc test; ^#^p<0.05, ^##^p<0.01, ^###^p<0.001 denotes significance versus controls; *p<0.05, **p<0.01, ***p<0.001 denotes significance versus MCD mice.

Furthermore, we found that treatment with ATX inhibitor Cpd17 ameliorated hepatic inflammation and steatosis, as examined by H&E and Oil-red-O staining respectively **(Fig. 7d-e)**. Additionally, the MCD-diet feeding resulted in increased intrahepatic protein expression of F4/80 (macrophage marker) and collagen I, compared to controls, as examined by immunostaining **(Fig. 7d-e)**. Strikingly, *in vivo* treatment with our ATX inhibitor led to a significantly decreased protein expression of both collagen I and F4/80, as shown in the microscopic images and quantitative staining analysis.

Moreover, total collagen content in the livers was evaluated through hydroxyproline assay that further confirmed an increased total collagen I concentration in MCD-livers, which was decreased in Cpd17 treated mice **(Fig. 7f)**.

Additionally, quantitative PCR revealed a decrease upon Cpd17 treatment in inflammatory markers F4/80, CCL2 and intracellular cell adhesion molecule (iCAM1), HSC specific marker α-SMA and liver steatosis marker CCAAT-enhancer-binding proteins (C/EBP)^46^ **(Supplementary Fig. 4)**.

### Cpd17 differentially regulate downstream signaling *in vivo*

Next, we investigated the mechanism of Cpd17 *in vivo* in both MCD-diet- and CCl4-induced liver disease models. In line with our *in vitro* analysis, we analyzed ERK, AKT and RhoA signaling pathways. It is important to note that these signaling pathways are involved in several cellular processes independent of ATX signaling, and analysis of the *in vivo* results can be challenging. Cpd17 significantly decreased ERK activation in both animal models (**Supplementary Fig. 5 and 6**), consistent with the *in vitro* results (Fig. 5c), but did not result in significant changes in pAKT activation *in vivo* (**Supplementary Fig. 5)**. Finally, the levels of phospho-Myosin Light Chain 2 (pMLC2) to monitor RhoA pathway activation, showed an increase in MCD mice, which was reduced to control levels following Cdp17 treatment; however, both changes were marginally not significant (**Supplementary Fig. 5)**. These results confirm that Cpd17 affects *in vivo* the same disease-relevant pathways as it affects in vitro and its mode of action.

## Discussion

The ATX/LPA signaling axis has previously shown to be involved with the initiation and progression of liver diseases^26-31^. Here, we show significant upregulation of ATX gene and protein expression in human liver NASH and cirrhosis patients, which we recapitulated in CCl_4_-induced liver injury and MCD diet-induced NASH mouse models. Having confirmed and strengthened the connection between ATX and liver disease, we showed that a (yet unexplored) type IV ATX inhibitor Cpd17, compared with previously reported type I ATX inhibitor PF8380, possess stronger inhibitory effects in a variety of *in vitro* assays in hepatocytes, pro-inflammatory macrophages and activated HSCs. We found that ATX inhibition by Cpd17 attenuated TGFβ-induced expression of fibrotic markers in human HSCs, and LPS/IFNγ-stimulated expression of inflammatory markers in both mouse and human macrophages. We also showed a decrease in collagen I contraction of HSCs upon ATX inhibition, a measure of functional cellular activation^47^, and decreased migration of activated HSCs and macrophages. LPA via Gα(12/13) activates RhoA kinase pathway by which it regulates HSCs morphology by organizing the actin cytoskeleton thereby HSCs-ECM interaction and HSCs contraction thus fibrotic phenotypes^45^. While examining the effect of Cpd17 and PF8380 on ATX/LPA-related downstream signaling, we evidenced that they notably affected the RhoA pathway both *in vitro* and *in vivo*. We also showed broader effects through the MAPK/ERK and AKT/PKB pathways, and that these signals go through LPA receptors. We thus establish that the mechanisms underlying the ATX inhibition, using a type IV inhibitor, have an effect on downstream signaling processes that influence inflammation and fibrogenesis.

The cell-based assays clearly demonstrated that Cpd17 has a notably better potential in ameliorating fibrotic phenotypes and downstream signaling responses, compared to PF8380, which had already shown effectiveness in attenuating CCl_4_-induced liver cirrhosis^32^. PF8380 is a so-called type I inhibitor of ATX, which affects LPC hydrolysis to the signaling LPA molecule. However, the type I inhibitor PF8380 does not occupy the allosteric tunnel of ATX that leads to activity modulation^48^ and has been suggested to be implicated in presenting LPA to its cognate GPCRs^49^. In contrast, Cpd17^36^ is a type IV inhibitor that prevents substrate binding and also occupies the ATX tunnel. Cpd17 is structurally (but not chemically) related to GLPG1690^24^, which has shown promise in treating IPF^50^.

Based on our *in vitro* results, we further evaluated ATX inhibition by Cpd17 *in vivo* in acute liver injury and NASH mouse models. We used the CCl_4_-induced liver fibrosis model^51^ and the MCD-diet-induced NASH mouse model^52^ which showed characteristic hepatocytes damage and/or lipid accumulation, accompanied by intra-hepatic inflammation and fibrosis, as reported previously^53^. Cpd17 has an effect both on MCD diet-induced hepatic steatosis, and in the CCl_4_-induced liver injury model. These results are in accordance with the previous study in which adipose-specific ATX deficiency exhibited reduced lipid accumulation and hepatic steatosis associated with high-fat diet, while ATX overexpression was found to aggravate steatosis^54^. Interestingly, previous studies with the potent Ex_31 inhibitor had no effect on CCl_4_-induced liver fibrosis and choline-deficient amino acid-defined diet-induced liver injury^34^. PAT-505, a type III ATX inhibitor, showed reduced fibrosis development in a high-fat diet-induced NASH mouse model with no significant effect on steatosis, inflammation, or hepatocyte ballooning^33^. Moreover, inhibition by PF8380 in CCl_4_-induced liver disease models has shown to reduce plasma ATX activity and liver LPA levels by approximately 50%, and attenuate fibrosis-based on histopathological scoring and collagen deposition in the liver^32^. Based on our results, ATX inhibition by Cpd17 showed protection against CCl_4_-induced acute liver injury, and decreased steatosis, inflammation, and fibrosis in the NASH mouse model.

The analysis of the signaling pathways examined *in vitro* were largely consistent with *in vivo*. Cdp17 significantly decreased pERK signaling in both animal models, and also decreased the pMLC marker of RhoA signaling back to physiological levels, albeit the latter change was marginally not statistically significant (p=0.07). Curiously, pAKT levels showed contradicting results upon Cpd17 treatment *in vitro* (decrease) and *in vivo* (increase). However, pAKT/AKT levels are differentially regulated in different cell types during liver injury and regeneration. On one hand, AKT activation improves hepatic regeneration, and is involved in restricting pro-inflammatory and promoting anti-inflammatory responses via negative regulation of TLR and NF-κB signaling in macrophages^55^; on the other hand, PI3K/AKT pathway blockade in HSCs has been shown to suppress fibrotic responses by inhibiting HSCs proliferation and collagen synthesis^56^. Interestingly, the ERK-AKT axis has been suggested to act as a switch regulating liver regeneration or fibrosis via liver sinusoidal endothelial cells (LSECs)-HSCs communication^57^. Thus, the expected effect of Cpd17 through LPA-mediated signaling, is likely masked by the effect of other relevant regulatory pathways in the MAP/AKT signaling pathway which affect differentially different cell types in the liver.

The excellent *in vitro* and *in vivo* efficacy of Cpd17 in liver disease models, together with GLPG1690 efficacy in treating IPF, argue that Type IV compounds are very promising agents in treating fibrotic diseases and possibly other pathologies related to the LPA/ATX axis. It must be noted, though, that the phase III Isabella trial for GLPG1690 was terminated due to adverse effects. Detailed results from the Isabella trial have not been published to this moment and it is thus hard to speculate why it failed. If that was due to the off-target effects of the specific compound, this suggests a need to assess alternative chemical scaffolds like Cpd17.

Based on the results in this study, we have clearly demonstrated the role of the ATX-LPA axis in NASH and liver cirrhosis. We have also shown that inhibition of ATX, using a type IV inhibitor Cpd17, ameliorates fibrosis, inflammation, and steatosis in HSCs and macrophages *in vitro*, as well as in NASH and liver cirrhosis mouse models. We also note, that while Cpd17 clearly effects phenotypes relevant to fibrosis in liver cells, it cannot be excluded that its therapeutic potential *in vivo*, might be related to an interplay with the immune system, and specifically with the infiltration of T-cells, as recently shown in a tumor model^12^. Regardless, type IV ATX inhibitors have clear promise in non-alcoholic steatohepatitis and liver cirrhosis. Further, our results also suggest it worth exploring the role of the ATX/LPA axis and the effect of its inhibition in models of liver cancer, with potential therapeutic opportunities for HCC.

## Acknowledgements

This research was funded by the Oncode Institute, the Netherlands Cancer Institute, and the University of Twente.

## References

1. Asrani, S.K., Devarbhavi, H., Eaton, J. & Kamath, P.S. Burden of liver diseases in the world. Journal of hepatology 70, 151–171 (2019).

2. Bernal, W., Auzinger, G., Dhawan, A. & Wendon, J. Acute liver failure. Lancet 376, 190–201 (2010).

3. Marcellin, P. & Kutala, B.K. Liver diseases: A major, neglected global public health problem requiring urgent actions and large-scale screening. Liver International 38, 2–6 (2018).

4. Younossi, Z. et al. Global burden of NAFLD and NASH: trends, predictions, risk factors and prevention. Nature reviews. Gastroenterology & hepatology 15, 11–20 (2018).

5. Byrne, C.D. & Targher, G. NAFLD: A multisystem disease. Journal of hepatology 62, S47–S64 (2015).

6. Kazankov, K. et al. The role of macrophages in nonalcoholic fatty liver disease and nonalcoholic steatohepatitis. Nature Reviews Gastroenterology & Hepatology 16, 145–159 (2019).

7. Francque, S.M., van der Graaff, D. & Kwanten, W.J. Non-alcoholic fatty liver disease and cardiovascular risk: Pathophysiological mechanisms and implications. Journal of hepatology 65, 425–443 (2016).

8. Anstee, Q.M., Reeves, H.L., Kotsiliti, E., Govaere, O. & Heikenwalder, M. From NASH to HCC: current concepts and future challenges. Nature reviews. Gastroenterology & hepatology 16, 411–428 (2019).

9. Konerman, M.A., Jones, J.C. & Harrison, S.A. Pharmacotherapy for NASH: Current and emerging. Journal of hepatology 68, 362–375 (2018).

10. Choi, J.W. et al. LPA receptors: subtypes and biological actions. Annual review of pharmacology and toxicology 50, 157–186 (2010).

11. Balupuri, A. et al. Discovery and optimization of ATX inhibitors via modeling, synthesis and biological evaluation. European Journal of Medicinal Chemistry 148, 397–409 (2018).

12. Matas-Rico, E. et al. Autotaxin impedes anti-tumor immunity by suppressing chemotaxis and tumor infiltration of CD8^+^ T cells. bioRxiv, 2020.2002.2026.966291 (2021).

13. van Meeteren, L.A. & Moolenaar, W.H. Regulation and biological activities of the autotaxin-LPA axis. Progress in lipid research 46, 145–160 (2007).

14. Kano, K., Aikawa, S., Hashimoto, T. & Aoki, J. Lysophosphatidic acid as a lipid mediator with multiple biological actions. The Journal of Biochemistry 157, 81–89 (2014).

15. Tigyi, G. & Parrill, A.L. Molecular mechanisms of lysophosphatidic acid action. Progress in lipid research 42, 498–526 (2003).

16. Sheng, X., Yung, Y.C., Chen, A. & Chun, J. Lysophosphatidic acid signalling in development. Development (Cambridge, England) 142, 1390–1395 (2015).

17. Mutoh, T., Rivera, R. & Chun, J. Insights into the pharmacological relevance of lysophospholipid receptors. British journal of pharmacology 165, 829–844 (2012).

18. Teo, S.T., Yung, Y.C., Herr, D.R. & Chun, J. Lysophosphatidic acid in vascular development and disease. IUBMB Life 61, 791–799 (2009).

19. Yung, Y.C., Stoddard, N.C. & Chun, J. LPA receptor signaling: pharmacology, physiology, and pathophysiology. J Lipid Res 55, 1192–1214 (2014).

20. Borza, R., Salgado-Polo, F., Moolenaar, W.H. & Perrakis, A. Structure and function of the ecto-nucleotide pyrophosphatase/phosphodiesterase (ENPP) family: Tidying up diversity. The Journal of biological chemistry 298, 101526 (2022).

21. Moolenaar, W.H. & Perrakis, A. Insights into autotaxin: how to produce and present a lipid mediator. Nature Reviews Molecular Cell Biology 12, 674 (2011).

22. Hausmann, J. et al. Structural basis of substrate discrimination and integrin binding by autotaxin. Nature structural & molecular biology 18, 198–204 (2011).

23. Geraldo, L.H.M. et al. Role of lysophosphatidic acid and its receptors in health and disease: novel therapeutic strategies. Signal Transduction and Targeted Therapy 6, 45 (2021).

24. Salgado-Polo, F. & Perrakis, A. The Structural Binding Mode of the Four Autotaxin Inhibitor Types that Differentially Affect Catalytic and Non-Catalytic Functions. Cancers 11, 1577 (2019).

25. Zulfikar, S., Mulholland, S., Adamali, H. & Barratt, S.L. Inhibitors of the Autotaxin-Lysophosphatidic Acid Axis and Their Potential in the Treatment of Interstitial Lung Disease: Current Perspectives. Clin Pharmacol 12, 97–108 (2020).

26. Kondo, M. et al. Increased serum autotaxin levels in hepatocellular carcinoma patients were caused by background liver fibrosis but not by carcinoma. Clinica chimica acta; international journal of clinical chemistry 433, 128–134 (2014).

27. Nakagawa, H. et al. Autotaxin as a novel serum marker of liver fibrosis. Clinica chimica acta; international journal of clinical chemistry 412, 1201–1206 (2011).

28. Fujimori, N. et al. Serum autotaxin levels are correlated with hepatic fibrosis and ballooning in patients with non-alcoholic fatty liver disease. World journal of gastroenterology 24, 1239–1249 (2018).

29. Yamazaki, T. et al. Association of Serum Autotaxin Levels with Liver Fibrosis in Patients with Chronic Hepatitis C. Scientific reports 7, 46705–46705 (2017).

30. Ando, W. et al. Serum Autotaxin Concentrations Reflect Changes in Liver Stiffness and Fibrosis After Antiviral Therapy in Patients with Chronic Hepatitis C. Hepatology communications 2, 1111–1122 (2018).

31. Joshita, S. et al. Serum autotaxin is a useful liver fibrosis marker in patients with chronic hepatitis B virus infection. Hepatology research : the official journal of the Japan Society of Hepatology 48, 275–285 (2018).

32. Kaffe, E. et al. Hepatocyte autotaxin expression promotes liver fibrosis and cancer. Hepatology (Baltimore, Md.) 65, 1369–1383 (2017).

33. Bain, G. et al. Selective Inhibition of Autotaxin Is Efficacious in Mouse Models of Liver Fibrosis. The Journal of pharmacology and experimental therapeutics 360, 1–13 (2017).

34. Baader, M. et al. Characterization of the properties of a selective, orally bioavailable autotaxin inhibitor in preclinical models of advanced stages of liver fibrosis. British journal of pharmacology 175, 693–707 (2018).

35. Salgado-Polo, F. et al. Autotaxin facilitates selective LPA receptor signaling. bioRxiv, 2022.2004.2009.487723 (2022).

36. Keune, W.-J. et al. Rational Design of Autotaxin Inhibitors by Structural Evolution of Endogenous Modulators. Journal of Medicinal Chemistry 60, 2006–2017 (2017).

37. Kedziora, K.M. et al. Rapid Remodeling of Invadosomes by Gi-coupled Receptors: DISSECTING THE ROLE OF Rho GTPases. The Journal of biological chemistry 291, 4323–4333 (2016).

38. Akcora, B.O., Storm, G. & Bansal, R. Inhibition of canonical WNT signaling pathway by beta-catenin/CBP inhibitor ICG-001 ameliorates liver fibrosis in vivo through suppression of stromal CXCL12. Biochimica et biophysica acta. Molecular basis of disease 1864, 804–818 (2018).

39. Joshi-Barve, S. et al. Palmitic acid induces production of proinflammatory cytokine interleukin-8 from hepatocytes. Hepatology (Baltimore, Md.) 46, 823–830 (2007).

40. Seki, E. & Schwabe, R.F. Hepatic inflammation and fibrosis: functional links and key pathways. Hepatology (Baltimore, Md.) 61, 1066–1079 (2015).

41. Zhang, C.Y., Yuan, W.G., He, P., Lei, J.H. & Wang, C.X. Liver fibrosis and hepatic stellate cells: Etiology, pathological hallmarks and therapeutic targets. World J Gastroenterol 22, 10512–10522 (2016).

42. He, L., Yuan, H., Liang, J., Hong, J. & Qu, C. Expression of hepatic stellate cell activation-related genes in HBV-, HCV-, and nonalcoholic fatty liver disease-associated fibrosis. PloS one 15, e0233702 (2020).

43. Gurvich, O.L. et al. Transcriptomics uncovers substantial variability associated with alterations in manufacturing processes of macrophage cell therapy products. Sci Rep 10, 14049 (2020).

44. Wang, Z. et al. Autotaxin stimulates LPA2 receptor in macrophages and exacerbates dextran sulfate sodium-induced acute colitis. Journal of molecular medicine (Berlin, Germany) 98, 1781–1794 (2020).

45. Yanase, M. et al. Lysophosphatidic acid enhances collagen gel contraction by hepatic stellate cells: association with rho-kinase. Biochem Biophys Res Commun 277, 72–78 (2000).

46. Rahman, S.M. et al. CCAAT/enhancing binding protein β deletion in mice attenuates inflammation, endoplasmic reticulum stress, and lipid accumulation in diet-induced nonalcoholic steatohepatitis. Hepatology (Baltimore, Md.) 45, 1108–1117 (2007).

47. Corin, K.A. & Gibson, L.J. Cell contraction forces in scaffolds with varying pore size and cell density. Biomaterials 31, 4835–4845 (2010).

48. Salgado-Polo, F. et al. Lysophosphatidic acid produced by autotaxin acts as an allosteric modulator of its catalytic efficiency. The Journal of biological chemistry 293, 14312–14327 (2018).

49. Nishimasu, H. et al. Crystal structure of autotaxin and insight into GPCR activation by lipid mediators. Nature structural & molecular biology 18, 205–212 (2011).

50. Desroy, N. et al. Discovery of 2-[[2-Ethyl-6-[4-[2-(3-hydroxyazetidin-1-yl)-2-oxoethyl]piperazin-1-yl]-8-methyli midazo[1,2-a]pyridin-3-yl]methylamino]-4-(4-fluorophenyl)thiazole-5-carbonitrile (GLPG1690), a First-in-Class Autotaxin Inhibitor Undergoing Clinical Evaluation for the Treatment of Idiopathic Pulmonary Fibrosis. J Med Chem 60, 3580–3590 (2017).

51. Scholten, D., Trebicka, J., Liedtke, C. & Weiskirchen, R. The carbon tetrachloride model in mice. Lab Anim 49, 4–11 (2015).

52. Machado, M.V. et al. Mouse models of diet-induced nonalcoholic steatohepatitis reproduce the heterogeneity of the human disease. PloS one 10, e0127991 (2015).

53. Nevzorova, Y.A., Boyer-Diaz, Z., Cubero, F.J. & Gracia-Sancho, J. Animal models for liver disease - A practical approach for translational research. Journal of hepatology 73, 423–440 (2020).

54. Brandon, J.A. et al. Adipose-derived autotaxin regulates inflammation and steatosis associated with diet-induced obesity. PloS one 14, e0208099 (2019).

55. Vergadi, E., Ieronymaki, E., Lyroni, K., Vaporidi, K. & Tsatsanis, C. Akt Signaling Pathway in Macrophage Activation and M1/M2 Polarization. J Immunol 198, 1006–1014 (2017).

56. Son, M.K. et al. HS-173, a novel PI3K inhibitor, attenuates the activation of hepatic stellate cells in liver fibrosis. Sci Rep 3, 3470 (2013).

57. Lao, Y. et al. Targeting Endothelial Erk1/2-Akt Axis as a Regeneration Strategy to Bypass Fibrosis during Chronic Liver Injury in Mice. Mol Ther 26, 2779–2797 (2018).

